# A non-adaptive demographic mechanism for genome expansion in *Streptomyces*

**DOI:** 10.1101/2021.01.09.426074

**Authors:** Mallory J Choudoir, Marko J Järvenpää, Pekka Marttinen, Daniel H Buckley

**Affiliations:** School of Integrative Plant Science, Cornell University, Ithaca, NY, USA; Department of Computer Science, Helsinki Institute for Information Technology HIIT, Aalto University, Espoo, Finland

## Abstract

The evolution of microbial genome size is driven by gene acquisition and loss events that occur at scales from individual genomes to entire pangenomes. The equilibrium between gene gain and loss is shaped by evolutionary forces, including selection and drift, which are in turn influenced by population demographics. There is a well-known bias towards deletion in microbial genomes, which promotes genome streamlining. Less well described are mechanisms that promote genome expansion, giving rise to the many microbes, such as *Streptomyces*, that have unusually large genomes. We find evidence of genome expansion in *Streptomyces* sister-taxa, and we hypothesize that a recent demographic range expansion drove increases in genome size through a non-adaptive mechanism. These *Streptomyces* sister-taxa, NDR (northern-derived) and SDR (southern-derived), represent recently diverged lineages that occupy distinct geographic ranges. Relative to SDR genomes, NDR genomes are larger, have more genes, and their genomes are enriched in intermediate frequency genes. We also find evidence of relaxed selection in NDR genomes relative to SDR genomes. We hypothesize that geographic range expansion, coupled with relaxed selection, facilitated the introgression of non-adaptive horizontally acquired genes, which accumulated at intermediate frequencies through a mechanism known as genome surfing. We show that similar patterns of pangenome structure and genome expansion occur in a simulation that models the effects of population expansion on genome dynamics. We show that non-adaptive evolutionary phenomena can explain expansion of microbial genome size, and suggest that this mechanism might explain why some bacteria with large genomes can be found in soil.

## Introduction

Microbial genomes are extraordinarily dynamic. Genome size varies considerably, and gene content in strains of the same species can differ dramatically, giving rise to the pangenome. The pangenome concept has transformed our understanding of evolutionary processes in diverse taxa [1–4]. The pangenome is the entire collection of genes in a microbial species, and is subdivided into core genes present in all strains, dispensable or accessory genes present in some strains, and strain-specific or unique genes [5, 6]. Rates of gene acquisition and gene loss determine the individual genome size, and consequently, pangenome composition is shaped by evolutionary mechanisms that alter gene frequencies in microbial populations [7–9].

Genome size varies by four orders of magnitude (10^4^–10^7^ kb) in eukaryotic organisms and two orders of magnitude in prokaryotic organisms (from less than 150 kb in certain endosymbionts to over 10 Mb for some free-living bacteria) [10]. Unlike eukaryotes, whose genomes contain large portions of non-coding DNA, prokaryotic gene content is directly related to genome size because bacterial and archaeal taxa have high coding density [11, 12]. While microbial genomes are constantly in flux, deletion rates are approximately three-fold greater than rates of gene acquisition [13]. Multiple factors contribute to the strong deletion bias in microbial genomes, including selection for efficiency, “use it or lose it” purging of nonessential genes, and genetic drift [14–17].

Because of this tendency towards deletion, microbial genome reduction has been examined in greater detail than genome expansion. For example, the evolutionary mechanisms driving genome reduction in obligate pathogens like *Rickettsia* and symbionts like *Buchnera* in aphids are well described [17, 18]. The transition from a free-living to a host-associated lifestyle involves substantial loss of superfluous genes, and generations of vertical transmission in small asexual populations leads to gene inactivation and deletion accelerated by genetic drift (16).

Alternatively, genome streamlining leads to reduction of both genome and cell size through selection for increased metabolic efficiency in free-living microbes with large populations [15]. Genome streamlining is historically associated with marine oligotrophic *Pelagibacter* [14, 19] but has more recently been described for soil-dwelling *Verrucomicrobia* [20].

Large genomes are frequent among terrestrial free-living microbes, and must be the product of evolutionary forces that drive genome expansion. A common, though relatively untested, hypothesis to explain large genomes is that high environmental heterogeneity (a characteristic of terrestrial habitats) selects for metabolic versatility afforded by gene gain, and thereby drives genome expansion [21, 22]. For example, massive gene acquisition and adaptation to alkaline conditions caused genome expansion in the myxobacterium *Sorangium cellulosum*, which at 15 Mb is one of the largest known bacterial genomes [23]. Mechanisms of gene gain include duplication or horizontal gene transfer (HGT), and large genomes are enriched in functional genes acquired from phylogenetically distant origins [24]. Much of the evolution of gene families can be attributed to HGT rather than duplication events [25, 26], and HGT is a major driver of genome expansion [27, 28]. While HGT-mediated gene acquisition occurs with great frequency, microbial genomes remain relatively small, and genome size tends to be fairly conserved within a species.

Gene frequencies at the population-level are governed by selection and drift, and these evolutionary forces determine whether a newly acquired gene will be purged from the pangenome or whether it will sweep to fixation. The strength of selection and drift varies inversely, and their relative contributions are determined by a gene’s selection coefficient and effective population size (N_e_) [29, 30]. Drift can exert large effects on populations with small N_e_, but these effects decline as N_e_ increases and selection intensifies. Our ability to disentangle the contributions of selection and drift to pangenome dynamics are complicated by the fact that it remains difficult to estimate microbial N_e_ [31, 32] and to delimit microbial population and species boundaries [33–35]. Another complication is that demographic models often include the simplifying expectation that N_e_ is invariable over time.

Rapid changes in population size are typical in the evolutionary histories of many microbial species, and fluctuations in N_e_ such as population bottlenecks or expansions can have profound impacts on contemporary patterns of genomic diversity. For example, the population structure for many pathogenic bacterial lineages is exemplified by episodes of rapid expansion of clonal complexes repeated across space and time [36–38]. Microbial population expansions can also be linked to ecological or geographical range expansions [39–42]. For instance, demographic expansion in the oral bacteria *Streptococcus mutans* coincides with the origin of human agricultural practices [41].

We find evidence for post-glacial range expansion in the genus *Streptomyces*, and these species exhibit several of the genetic characteristics described in plant and animal species whose biogeography was influenced by Pleistocene glaciation [43, 44]. By examining *Streptomyces* isolated from sites across North America, we observed genetic evidence for dispersal limitation, a latitudinal gradient in taxonomic richness, and a latitudinal gradient in genetic diversity [45, 46]. We also identified recently diverged sister-taxa comprising a more genetically diverse southern-derived (SDR) clade and a more homogenous northern-derived (NDR) clade, which occupied discrete geographic ranges spanning the boundary of glaciation [47]. We further observed larger genomes in the northern clade compared to the southern clade.

We hypothesize that genome expansion in NDR is a consequence of demographic change driven by post-Pleistocene range expansion. Here, we evaluate the effects of historical range expansion on lineage divergence, genome size, and pangenome structure, and assess these data in the context of the genome surfing hypothesis. Genome surfing is a non-adaptive mechanism which describes the introgression of horizontally acquired genes facilitated by relaxed selection and amplified by geographic expansion [48]. We hypothesize that range expansion, coupled with relaxed selection, dampened gene loss thereby facilitating an increase in non-adaptive, intermediate frequency genes in the NDR pangenome. We infer gene gain and loss dynamics by evaluating patterns of shared gene content between strains. We predict that the contribution of drift is greater in NDR compared to SDR, and determine the relative strength of selection by comparing genome-wide rates of amino acid substitution between clades. Finally, we evaluate our hypothesis by modeling population expansion under a regime of relaxed selection and ask whether these demographic conditions increase retention of horizontally acquired genes at intermediate frequencies, ultimately causing genome expansion.

## Material and Methods

### Streptomyces isolation and genomic DNA extraction

The strains in this study belong to a larger culture collection of *Streptomyces* isolated from surface soils (0–5 cm) spanning sites across the United States (see [45, 46]) (Table S1). To minimize the effects of environmental filtering in driving patterns of microbial diversity, we selected sample locations with similar ecologies including meadow, pasture, or native grasslands dominated by perennials and with moderately acidic to neutral soils (pH 6.0 ± 1.0, mean ± SD).

*Streptomyces* strains were isolated by plating air-dried soils on glycerol-arginine agar (pH 8.7) plus cycloheximide and Rose Bengal [49, 50] as previously described [51]. Genomic DNA was extracted with a standard phenol/chloroform/isoamyl alcohol protocol from 72 h liquid cultures grown at 30°C with shaking in yeast extract-malt extract medium (YEME) + 0.5% glycine [52].

### Genome sequencing, assembly, and annotation

Genome sequencing, assembly, and annotation is previously described (see [47]). Briefly, we used the Nextera DNA Library Preparation Kit (Illumina, San Diego, CA, USA) to prepare sequencing libraries. Genomes were sequenced on an Illumina HiSeq2500 instrument with paired-end reads (2 x 100 bp). Genomes were assembled with the A5 pipeline [53] and annotated with RAST [54]. This generated high quality draft genome assemblies with over 25X coverage and estimated completeness > 99% as assessed with CheckM [55]. We used ITEP and MCL clustering (inflation value = 2.0, cutoff = 0.04, maxbit score) [56] to identify orthologous protein-coding gene clusters (i.e., genes). Genome sequences are available through NCBI under BioProject PRJNA401484 accession numbers SAMN07606143–SAMN07606166.

### Phylogeny

Phylogenetic relationships were reconstructed from whole genome alignments. We used Mugsy [57] to generate multiple genome nucleotide alignments and trimAl v1.2 [58] for automatic trimming of poorly aligned regions. Maximum likelihood (ML) trees were built using the generalized time reversible nucleotide substitution model [59] with gamma distributed rate heterogeneity among sites (GTRGAMMA) in RAxML v7.3.0[60], and bootstrap support was determined following 20 ML searches with 100 inferences using the RAxML rapid bootstrapping algorithm [61]. Average nucleotide identity (ANI) was calculated from whole genome nucleotide alignments using mothur [62].

### Pangenome and population genetics analyses

The pangenome was determined from the gene content of 24 *Streptomyces* genomes (Table S2). Strains in this collection were initially chosen for whole genome sequencing based on their genetic similarity at house-keeping loci (see [46]). Subsequent analyses focused on recently diverged sister-taxa clades of 10 genomes each, the northern-derived (NDR) and southern-derived (SDR) lineages. Gene content patterns between strains and pangenome gene frequency distributions were determined from gene presence/absence data.

Gene-level attributes across gene pools were determined from the average of all nucleotide sequences within an orthologous protein-coding gene cluster (see above). GC content was calculated for each gene using the R package Biostrings [63]. Codon usage bias was calculated for each gene using the R package cordon [64]. Clade-level population genetic traits were evaluated using 2,778 single-copy genes conserved across all 24 genomes. For each core gene, nucleotide sequences were aligned using MAFFT v.7 [65], and Gblocks [66] removed poorly aligned positions. PAL2NAL [67] generated codon alignments, and SNAP [68] calculated intra-clade non-synonymous (K_A_) and synonymous (K_S_) substitution rates (values > 2 were filtered prior to plotting and statistical analysis).

### Demographic simulation

We assumed that the SDR pangenome approximates the gene frequency distribution of the last common ancestor of NDR and SDR. For the starting generation 0, we used the model from Marttinen *et al*. [69] to simulate a population of sequences and learn parameter values for rates of gene acquisition and deletion that produced the frequency distribution for SDR. To model range expansion demographics (i.e., severe bottleneck followed by exponential growth), we sampled 5 strains from generation 0 as the founding population for the subsequent generation, and simulated this for 100 generations. The simulated population had a growth rate of 5% per generation until a maximum of 100 individuals was reached. We varied the initial sizes of the founding population as well as the growth rate, and observed qualitatively similar results.

The model included gene acquisition events and deletion events similar to Marttinen *et al*. [69] but modified to allow for multiple changes. Instead of acquisitions/deletions happening independently, there were k=20 simultaneous acquisitions/deletions per strain per generation. The previous model [69] included a multiplicative fitness penalty of 0.99 for each gene exceeding a pre-specified genome size threshold. During the expansion, we relaxed the penalty for excess genes to 0.99^(current size/max size)^ allowing for genome size variation.

## Results

### Streptomyces sister-taxa

We sequenced the genomes of 20 *Streptomyces* strains isolated from ecologically similar grasslands sites across the United States (Table S1, Table S2). These genomes derive from sister-taxa comprising a northern-derived (NDR) and southern-derived clade (SDR), which originate from sites spanning the historical extent of glaciation (Figure S1, see [45]). These sister-taxa represent closely related but genetically distinct microbial species. Genomes within NDR share 97.8 ± 1.3% (mean ± SD) ANI and those within SDR share 97.6 ± 0.1% (mean ± SD) ANI, while inter-clade genomic ANI is 93.0 ± 0.14% (mean ± SD). An ANI of 93–96% is typically indicative of taxonomic species boundaries [70, 71]. For comparative purposes, we also sequenced the genomes of four strains that co-localized with the sister-taxa. The closest taxonomic neighbor to these 24 strains is *Streptomyces griseus* subsp. *griseus* NBRC 13350, although all strains share < 95% ANI with this type strain (Figure S1).

### Genomic attributes and gene content

NDR genomes are larger (8.70 ± 0.23 Mb, mean ± SD) than SDR genomes (7.87 ± 0.19 Mb, mean ± SD), and this difference is significant (Mann Whitney U test; *P* < 0.0001) (Figure 1a). NDR genomes also have also have more orthologous protein-coding gene clusters (hereby referred to as genes) (7,775 ± 196 genes, mean ± SD) than SDR genomes (7,093 ± 205 genes, mean ± SD), and this difference is also significant (Mann Whitney U test; *P* < 0.0001) (Figure 1b). As expected, there is a strong positive correlation between genome size and gene content (R^2^ = 0.95, *P* < 0.0001), but coding density did not differ between clades (Figure S2). SDR genomes are more genetically diverse than NDR. Nucleotide diversity (n) across conserved, single-copy core genes is greater in SDR than NDR, and this difference is significant (Mann Whitney U test; *P* < 0.0001) (see [47]). Finally, NDR genomes have slightly lower genome-wide GC content (71.50 ± 0.087%, mean ± SD) than SDR genomes (71.62 ± 0.11%, mean ± SD), and this difference is significant (Mann Whitney U test; *P* = 0.017) (Figure 1c). Shared gene content between strains correlates with genomic similarity as measured by ANI (NDR: R^2^ = 0.82, *P* < 0.0001; SDR: R^2^ = 0.64, *P* < 0.0001) (Figure 2). However, gene content varies more in NDR than in SDR, and there is a significant interaction between genomic similarity and clade with respect to gene content shared between strains (Table S3). This interaction comes from shared gene content between strains increasing more rapidly over recent phylogenetic timescales in NDR compared to SDR (Figure 3).

**Figure 1.**
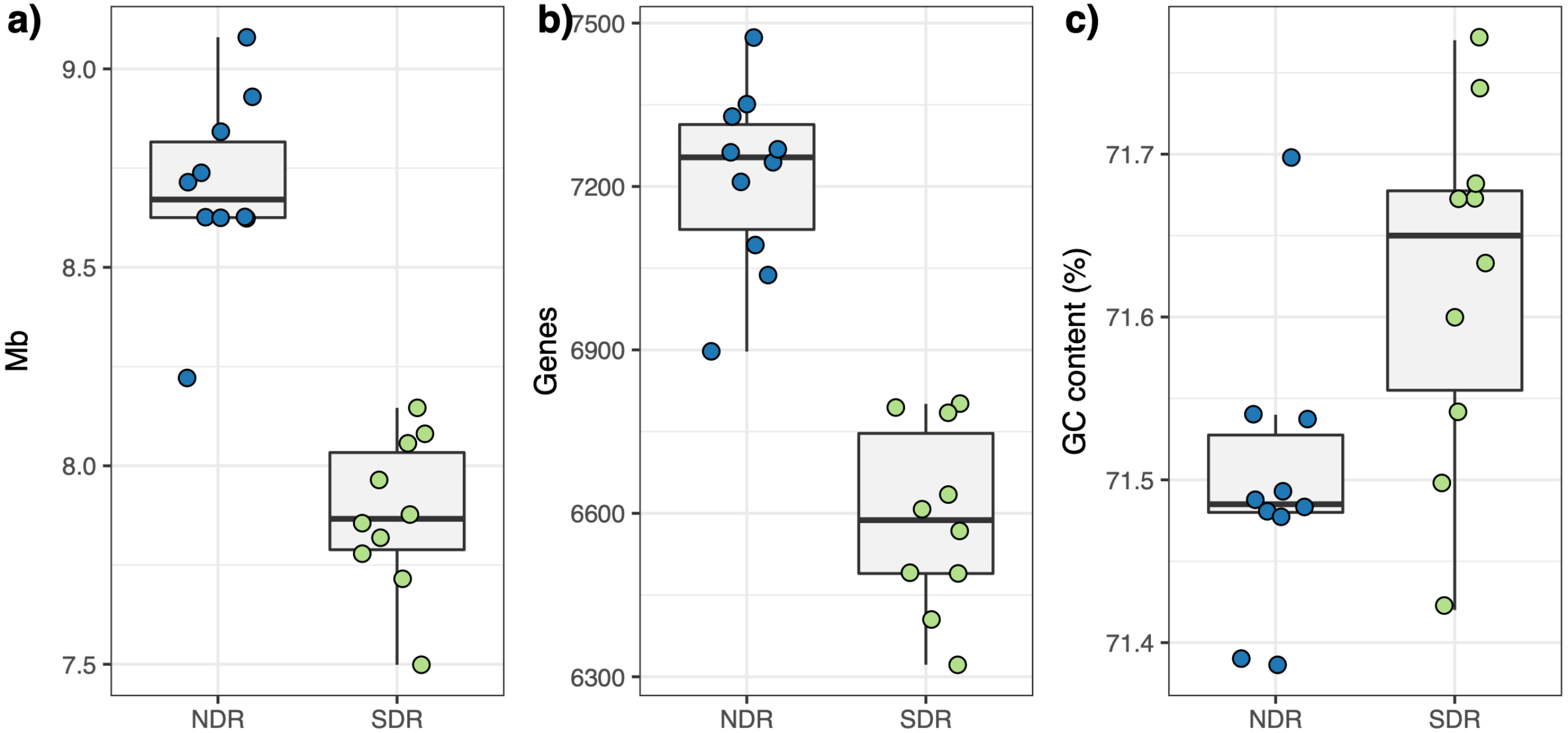
Genomic attributes of NDR and SDR sister-taxa. NDR genomes are larger, have more genes, and have lower GC content compared to SDR genomes. Plots show the distributions of genome size in Mb (a), number of genes (b), and genome-wide GC content (%) (c) for *Streptomyces* sister-taxa. Boxplots show the clade-level medians, interquartile ranges, and 1.5 times interquartile ranges. Colored circles illustrate the values for individual genomes belonging to the NDR clade (blue) or the SDR clade (green).

**Figure 2.**
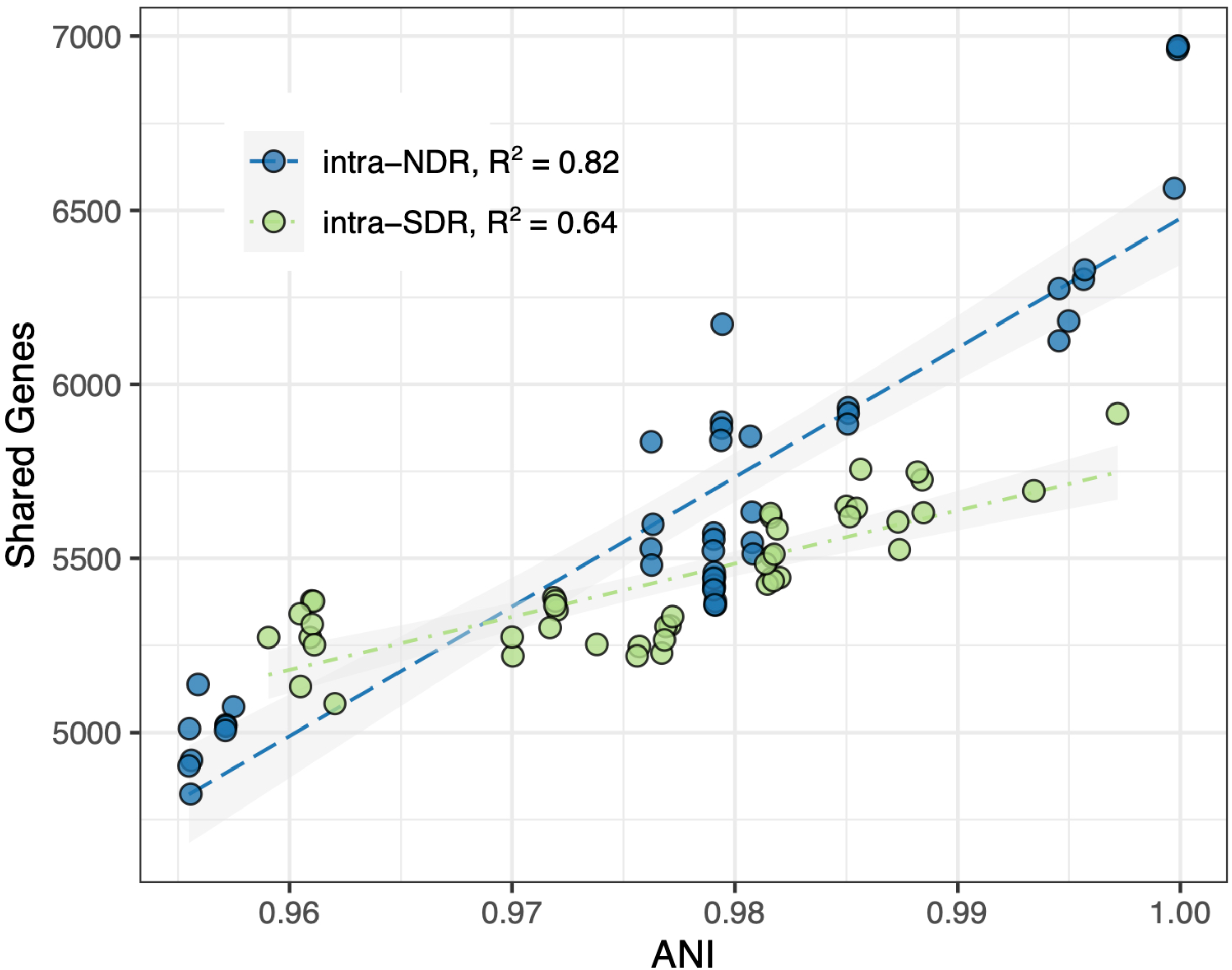
Genomic similarity versus shared gene content for NDR and SDR. Differences in shared gene content across increasing average nucleotide identity (ANI) are greater within the NDR clade compared to the SDR clade (Table S3). Circles show pairwise comparisons of the number of shared genes between two strains versus ANI and are colored by clade according to the legend. Dashed lines show linear regressions, and the shaded area is the 95% confidence interval.

**Figure 3.**
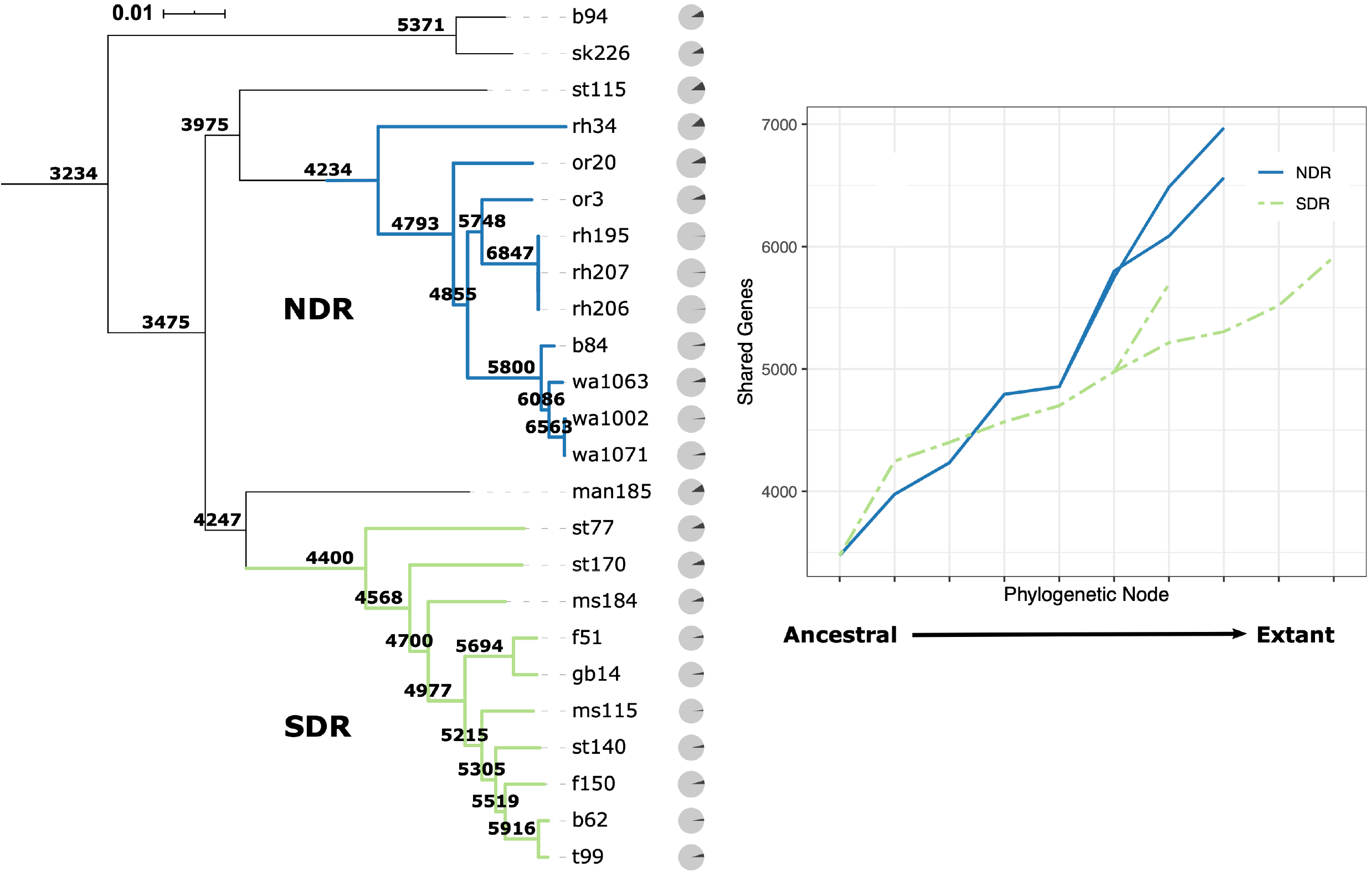
Presence/absence of genes across phylogeny. Gene content changes more rapidly across ancestral phylogenetic nodes for NDR genomes compared to SDR genomes. Tree is made from whole genome nucleotide alignments, and the scale bar shows nucleotide substitutions per site (see Figure S1). Branch colors reflect clade membership. Phylogenetic nodes are labeled with the number of genes conserved in all members of descendent nodes. Gray pie charts at tree tips show the portion of total genes per genome that are strain-specific (black slice). Right panel plots the differences in gene content across the phylogeny beginning at the shared ancestral node and ending with extant taxa at the terminal tips for NDR (blue-solid) and SDR (green-dashed) lineages. Multiple lines represent monophyletic lineages.

### Pangenome structure and dynamics

The 24 *Streptomyces* genomes (Table S2) contain 22,055 total orthologous protein-coding gene clusters (i.e., genes), and 42% (9,285 genes) are strain-specific. All 24 genomes share 3,234 (2,778 single-copy) genes, which represent 40–48% of the total gene content per strain. While NDR has a smaller core genome than SDR (4,234 and 4,400 genes, respectively), its pangenome is larger (13,681 genes in NDR versus 12,259 genes in SDR) and contains a greater number of clade-specific genes (5,647 genes unique to NDR versus 4,308 genes unique to SDR) (Figure 3, Figure 4, Table S4).

**Figure 4.**
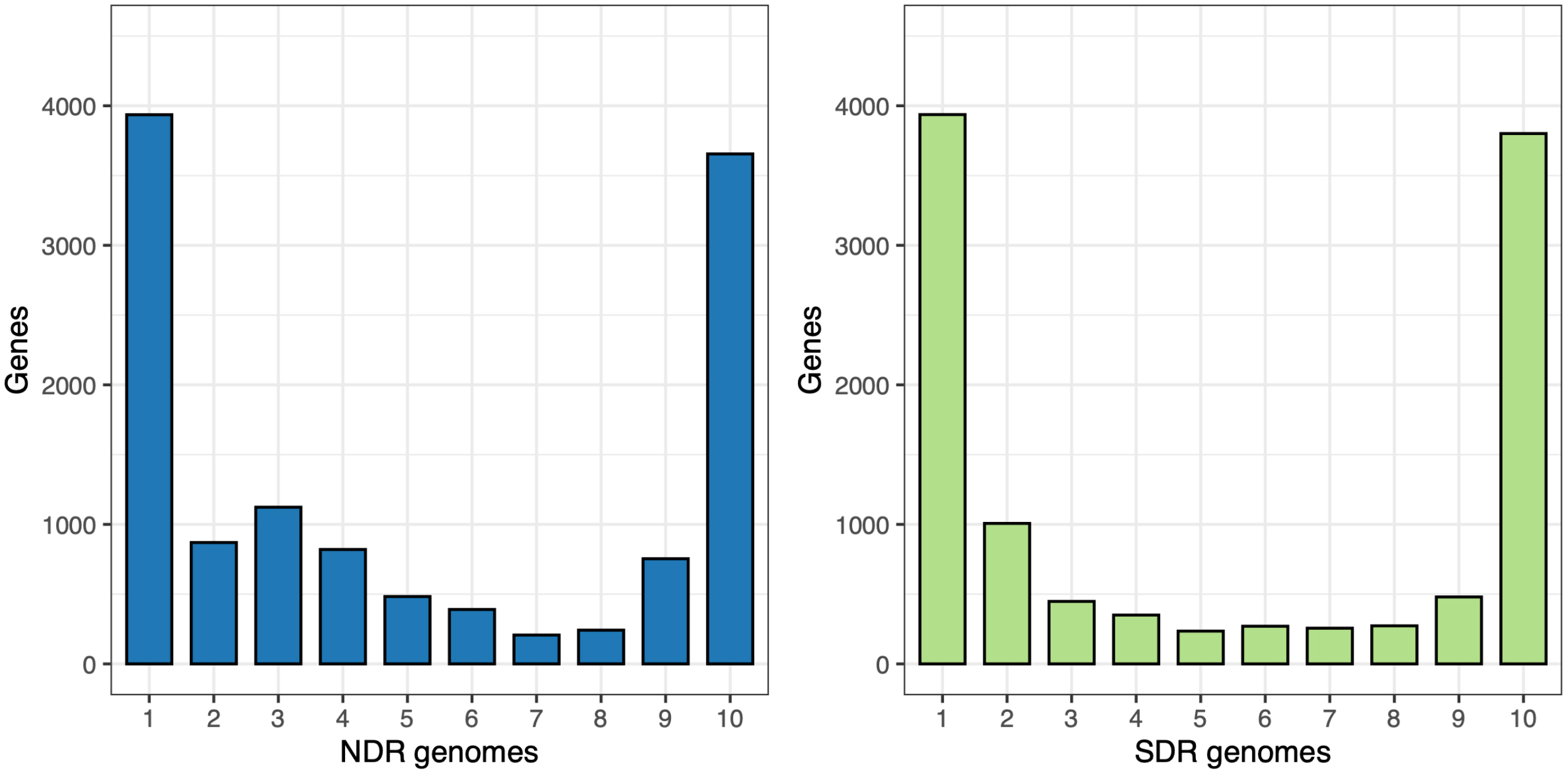
Pangenome gene frequency distributions. NDR genomes are enriched in intermediate frequency genes. Plots show the pangenome gene frequency distributions for NDR (left) and SDR (right). Bars show the population-level sums of genes present in 1–10 genomes. See Table S3 for raw values and proportions.

For most microbial species, pangenome frequency distributions are U-shaped, reflecting high proportions of both strain-specific genes and core genes [72]. While the pangenome structures of our *Streptomyces* sister-taxa generally conform to this shape, the NDR pangenome is enriched in intermediate frequency accessory genes relative to SDR (Figure 4). The proportion of intermediate-low frequency (i.e., present in 3–5 strains) accessory genes is higher in NDR than in SDR (19% of total genes for NDR versus 9.2% of total genes for SDR) (Table S4), and this difference is statistically significant (two proportion z-test; *P* < 0.0001). Conversely, the proportion of intermediate-high frequency (i.e., present in 6–8 strains) accessory genes is equivalent (6.9% of total genes for NDR versus 7.2% of total genes for SDR; two proportion z-test; *P* = 0.26) (Table S4).

Next, we determined if genes across different gene pools, binned according to their pangenome frequencies, differed in genetic attributes including per-gene GC content and codon usage bias. GC content differs between gene pools for both NDR and SDR pangenomes (ANOVA; *F*_3, 25932_ = 267.5, *P*-value < 0.0001) (Figure S3). In general, GC content is greater in high frequency and core genes compared to rare and intermediate frequency genes for both sister-taxa. Codon usage bias as measured by the effective number of codons (ENC) [73] also differs between gene pools for both NDR and SDR pangenomes (ANOVA; *F*_3, 21624_ = 1862.7, *P*-value < 0.0001) (Figure S4). Rare and intermediate frequency genes exhibit less overall codon bias compared to high frequency and core genes, which tend to use codons more preferentially.

### Historical population demography

Due to founder effects occurring at the edge of an expanding population, N_e_ is dramatically reduced during geographic range expansion [74]. Consequently, relaxed selection will accompany range expansion since the contribution of selection scales directly with N_e_. Based on the theory of neutral molecular evolution, which states that selection on synonymous sites is negligible [75], the ratio of non-synonymous to synonymous amino acid substitutions (K_A_/K_S_) reflects the relative strength of selection acting on a sequence. When assessed at the level of single-copy genes conserved between the sister taxa (2,444 genes), we observe that genome-wide K_A_/K_S_ tends to be higher in NDR than in SDR (Figure 5), and this difference is significant (Mann-Whitney U test; *P* < 0.0001). This result indicates that selection is weaker and genetic drift stronger in NDR relative to SDR.

**Figure 5.**
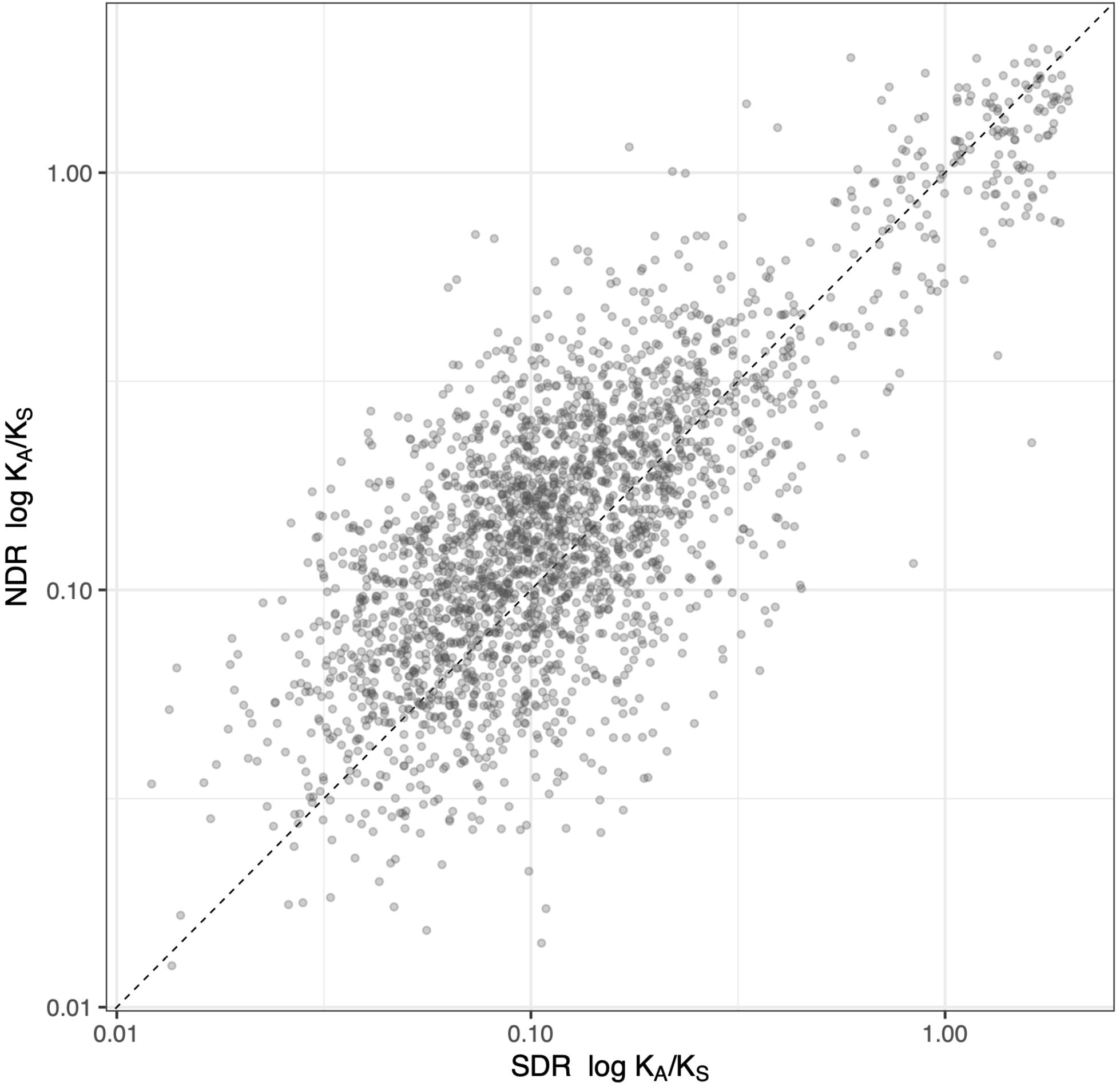
K_A_/K_S_ values between the NDR and SDR sister-taxa core genome. NDR core genes have, on average, greater rates of non-synonymous to synonymous amino acid substitutions compared to SDR core genes. Circles plot clade-level rates of non-synonymous to synonymous amino acid substitutions (K_A_/K_S_) for each of 2,444 single-copy core genes for NDR (y-axis) and SDR (x-axis). Axes are logarithmic scale. The black dashed line is a slope of 1, and points along this line are genes with equal K_A_/K_S_ mean values in both clades. K_A_/K_S_ is proportional to the relative strength of genetic drift and inversely proportional to the relative strength of selection.

We used a population model (modified from [69]) to determine whether demographic expansion could produce increased intermediate gene frequencies and result in genome expansion. We simulated gene gain and loss events in a population undergoing exponential growth over 100 generations, and determined changes in pangenome structure and genome size. To approximate relaxed selection during the population expansion, we imposed a fitness penalty for newly acquired genes that scaled inversely with population size. At the beginning of expansion, most genes were present at high frequencies due to strong founder effects (Figure S5, top and middle). During the expansion, we observed a transient enrichment of intermediate frequency genes within the pangenome (Figure S5, top and middle). Total gene content also increased during population expansion due to relaxed selection pressure when N_e_ was small, which allowed for the persistence of newly HGT-acquired genes. Genome size stabilized when N_e_ reached maximum size, and selection pressure balanced HGT-mediated gene gain with simultaneous gene loss (Figure S5, bottom).

## Discussion

We have hypothesized that the biogeography of our *Streptomyces* sister-taxa is explained by historical demographic change driven by geologic and climatic events that occurred in the late Pleistocene [46, 47]. Following the last glacial maxima, North American plant and animal species rapidly colonized glacial retreat zones, and the genetic consequences of post-glacial expansion are well documented and include northern-ranged populations with low diversity that established vast geographic extent [43, 44]. We hypothesize that the recent common ancestor of NDR and SDR inhabited southern glacial refugia prior to the last glacial maxima (LGM). Post glaciation, NDR dispersed northward and colonized the latitudinal range it occupies today (see [46]). We previously described patterns of gene flow, genomic diversity, and ecological adaptation in these sister-taxa, with both adaptive and non-adaptive processes likely reinforcing lineage divergence [47]. Here, we evaluate the outcomes of historical range expansion on sister-taxa pangenome structure and genome size.

Expanding populations experience repeated founder effects as individuals along the leading edge disperse and colonize new landscapes, creating spatial patterns of genetic diversity akin to genetic drift [74]. Allele surfing, or gene surfing, is a non-adaptive mechanism that propagates rare alleles along an expanding edge such that neutral, or even deleterious, variants ‘surf’ to higher frequencies than would be expected under population equilibrium [76–78]. When applied to expanding microbial populations, gene surfing can facilitate genome surfing, a neutral mechanism acting at the pangenome level that causes rare genes to surf to higher frequencies independent of natural selection [48]. Below, we outline how historical range expansion and genome surfing could give rise to genome expansion in *Streptomyces*.

Genome surfing is most likely to occur in microbial populations with intermediate levels of dispersal and in taxa capable of HGT. Bacteria in the genus *Streptomyces* are ubiquitous in soil and produce desiccation and starvation resistant spores which are easily disseminated [52], making them ideal for studying patterns of biogeography dependent on dispersal limitation. Rates of HGT in *Streptomyces* are among the highest estimated across a range of bacterial species [51, 79, 80]. In many instances, HGT events occurred in ancestral lineages creating patterns of shared genetic ancestry and reticulate evolution in many extant *Streptomyces* species [81]. We previously observed a distance decay relationship between sites up to 6,000 km apart, indicative of dispersal limitation at intermediate spatial scales that allows detection of geographic patterns of diversity across the sampled range [45, 46]. We also found evidence of restricted gene flow between the core genomes of NDR and SDR [47].

Since NDR and SDR sister-taxa share a recent common ancestor (Figure S1), they must also share a common ancestral genome size. Hence, differences in genome size accompanying lineage divergence resulted from either genome expansion in NDR or genome reduction in SDR. Given that changes in genome size are ultimately the result of gene gain and loss, we first evaluated differences in shared gene content between NDR and SDR strains. We find greater variability in shared gene content in NDR compared to SDR (Figure 2, Figure 3). This result suggests relative gene content stability for SDR and gene content instability for NDR, most notably in recent phylogenetic history (Figure 3). Likewise, the pangenome of NDR exceeds that of SDR by over 1,000 genes. Evidence suggests that during range expansion, founders at the expansion edge disperse into new habitats and acquire genes from local gene pools asymmetrically at unequal rates, and gene flow is almost exclusively from local to invading genomes [82]. These data are consistent with the observation that that NDR has a larger, more diverse, and more dynamic pangenome than SDR due to introgression from local gene pools.

Regardless of their origin, most novel horizontally-acquired genes are neutral or nearly neutral [83]. In most situations, selection will balance gene gain with gene deletion, and genome size will remain relatively constant.

Genetic diversity in individuals at the leading edge of an expanding population is dramatically reduced, and their genomes experience relaxed selection pressure due to consecutive population bottlenecks and low N_e_ [84]. We find that NDR has lower genetic diversity [47] and higher rates of K_A_/K_S_ across its core genome relative to SDR (Figure 5), which is consistent with the prediction that NDR has experienced a period of relaxed selection relative to SDR. A positive correlation is observed between GC content and selection pressure on microbial genomes [85, 86], and genome expansion in *Chlamydia* has been linked to relaxed selection resulting in a decrease in genome-wide GC content [87]. We likewise observe a decrease in genome-wide GC content in NDR relative to SDR (Figure 1). Relaxed selection pressure in NDR would mitigate the natural bias towards deletion and permit genes acquired by HGT to persist in the genome, regardless of their adaptive coefficient. Microbial sectoring that accompanies geographic range expansion [88] would then allow these newly acquired genes to accumulate at intermediate frequencies in the pangenome. The fact that NDR has larger overall genome size and that relative selection pressure is lower in NDR than SDR, is contrary to the predictions of the metabolic versatility hypothesis of large genomes.

We hypothesize that relaxed selection and drift caused genome expansion in NDR. While these same mechanisms are known to promote genome reduction in endosymbionts and obligate pathogens [17, 18], it is important to recognize that these outcomes are not contradictory (Figure 6). Genome size is regulated by rates of gene gain and loss, the selective coefficient for each gene in the genome, and the strength of selection. Endosymbionts and obligate intracellular pathogens have small population sizes and accordingly, relaxed selection and stronger drift. Relaxed selection pressure should lessen deletion bias. But under these conditions, host compensation for microbial gene function radically alters selective coefficients of core genes, thereby favoring genome reduction, and slightly deleterious mutations accumulate over time via Muller’s ratchet [89, 90]. In addition, rates of HGT from non-host sources are essentially zero, since there is little opportunity for endosymbionts to interact with other microbial cells, resulting in a one way track to genome erosion. In contrast, for free-living microbes relaxed selection pressure should bring about genome expansion by shifting the selective coefficients of accessory genes towards neutral. For example, genome expansion in *Chlamydia* was driven by relaxed selection, recombination, and introgression [87]. In this way small population size can favor genome erosion in endosymbionts, while also favoring genome expansion in free-living organisms (Figure 6). Meanwhile, free-living organisms that have large population sizes and high selection pressure will experience high rates of deletion that purge unnecessary genes in order to promote genome streamlining [14, 15].

**Figure 6.**
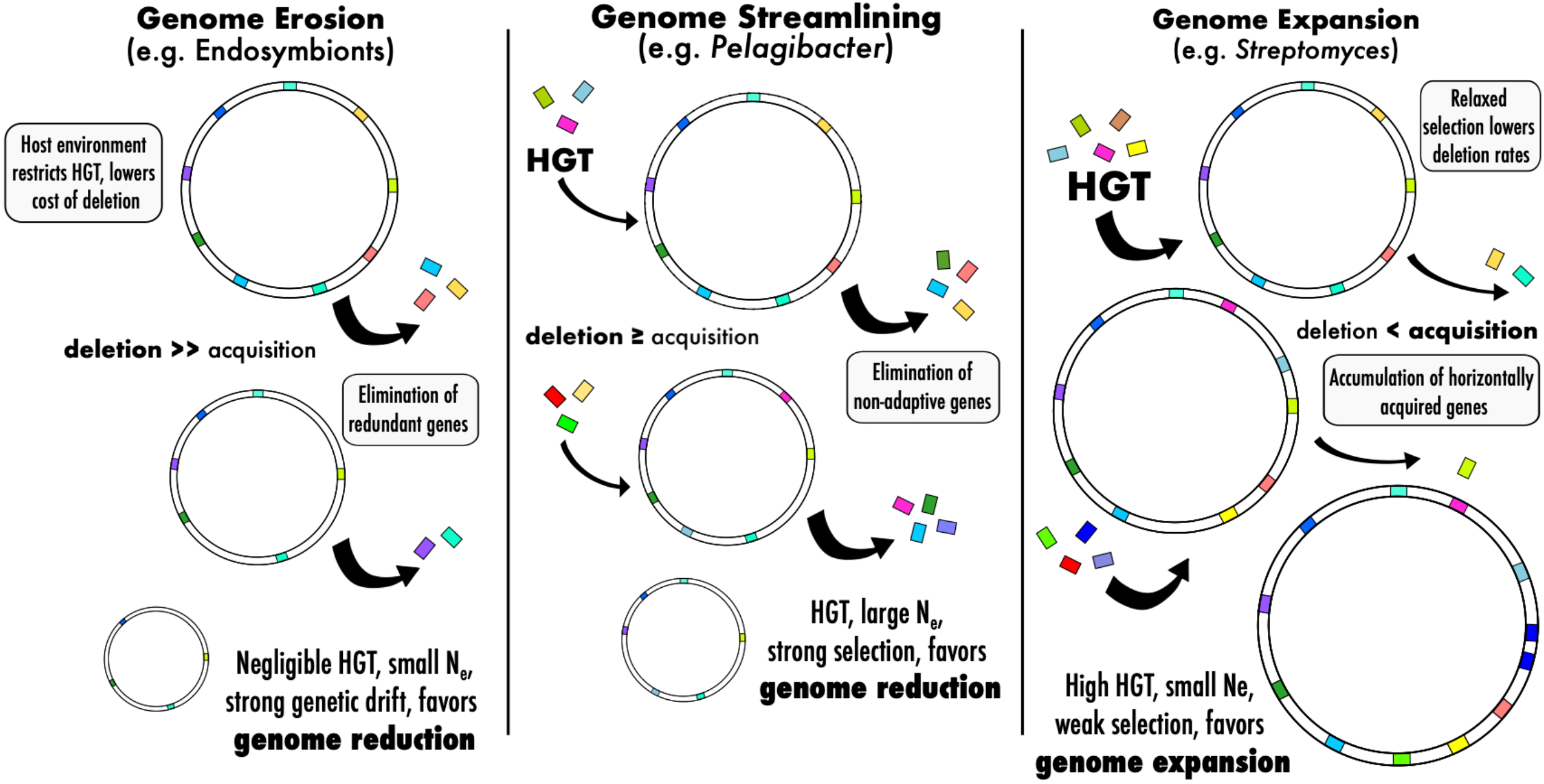
Conceptual overview of the evolutionary processes and demographic conditions that support changes in genome size. Genome erosion (left) in endosymbionts is the result of small N_e_ and strong genetic drift, with host compensation lowering costs of deletion while restricting gene flow (HGT). Genome streamlining (middle) in free-living microbes with large populations like *Pelagibacter* involves strong selection and elimination of non-adaptive genes. Genome expansion (right) in *Streptomyces* is facilitated by high rates of HGT and relaxed selection, allowing for the accumulation of non-adaptive genes and ultimately larger genomes.

Newly acquired genes tend to occur at low frequency in a population unless they provide an adaptive benefit [91], while adaptive genes will increase rapidly in frequency to join the core genome. These dynamics are believed to explain the characteristic U-shape of pangenome gene frequency distributions [72, 92]. Deviations from U-shape expectations, including increased intermediate frequency genes, can result from changes in selection coefficients of genes or under conditions where HGT exceeds deletion rates [93]. Alternatively, negative frequency dependent selection can cause highly beneficial genes to occur at low and intermediate frequencies [94, 95]. A large portion of rare genes in microbial pangenomes are hypothetical proteins or genes of unknown function acquired through HGT [96, 97]. For both NDR and SDR, approximately 60% of unique-rare genes (i.e., present in 1–2 strains) are annotated as hypothetical proteins. Nearly half of the 2,596 genes in NDR’s intermediate-low frequency gene pool (i.e., present in 3–5 strains) are also hypothetical genes. Furthermore, intermediate-low frequency genes are similar to rare frequency genes in regards to GC content (Figure S3) and codon usage (Figure S4). These data are consistent with our hypotheses that NDR intermediate frequency genes represent evolutionarily recent HGT-gene acquisitions, which increased in frequency as a result of genome surfing.

HGT-mediated genome expansion supplies a reservoir of novel genetic material for the evolution of gene families [25, 26], biosynthetic pathways [98], and formation of new metabolic networks [99]. Hence, the metabolic versatility of large genomes might be a classic example of an evolutionary spandrel [100], an adaptive trait associated with large genomes that originated not because of selection for versatility, but rather because the acquisition of diverse metabolic pathways is a byproduct of non-adaptive evolutionary process that cause genome expansion.

We show that pangenome analysis of *Streptomyces* sister-taxa verifies several predictions of the hypothesis that genome expansion within this clade was enabled by non-adaptive evolutionary processes, most likely driven by late Pleistocene demography. We hypothesize that small effective population size and relaxed selection, a consequence of geographic range expansion, allowed for genes newly acquired by HGT to increase in frequency within the NDR pangenome as a result of genome surfing. Further amplifying this effect is introgression of genes from local gene pools encountered following dispersal into new environments. Non-adaptive genome expansion is inherently a non-equilibrium process driven by a transient period of relaxed selection, and population stabilization will re-impose selection pressures that favor deletion. At this point, intermediate frequency genes will either be lost to deletion or fixed if they provide adaptive benefits, and these processes will shift the pangenome structure back to U-shaped expectations. These insights highlight the importance of considering population demography and the profound influence of historical contingency on contemporary patterns of microbial genome diversity.

## Data Availability

*Streptomyces* genome sequences are available through NCBI under BioProject PRJNA401484 accession numbers SAMN07606143–SAMN07606166.

## Acknowledgements

This work was supported by the National Science Foundation under Grant No. DEB-1456821 awarded to Daniel H Buckley.

## Supplementary Information for

**Table S1.**
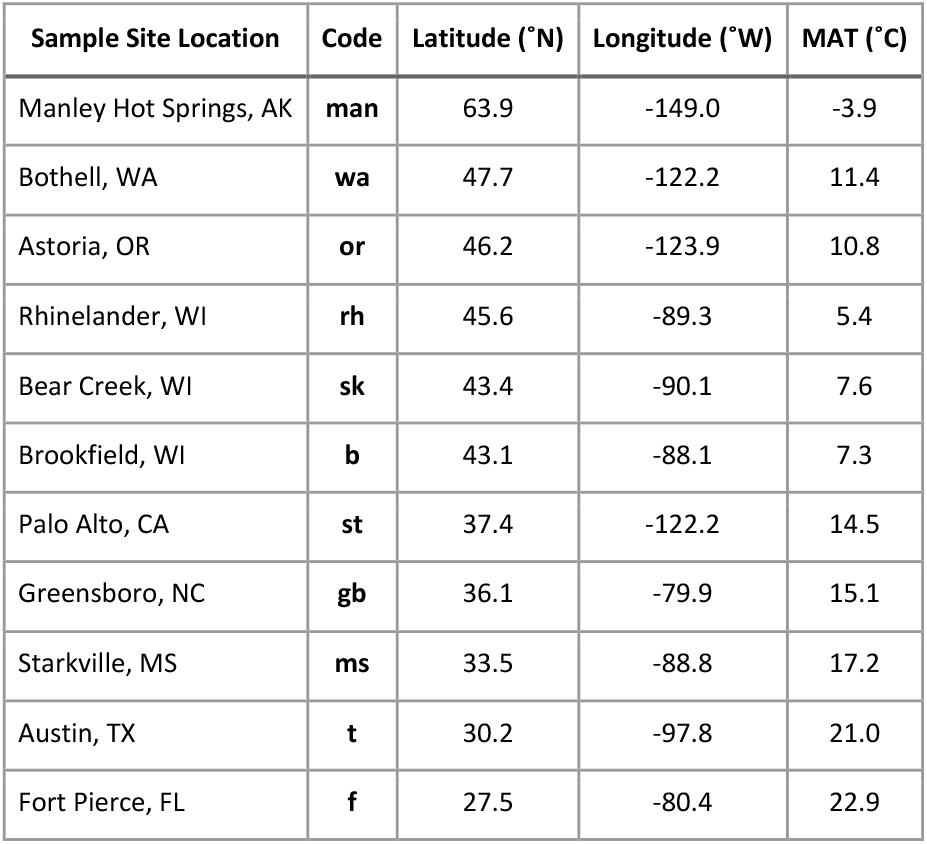
*Streptomyces* strains were isolated from 11 grassland sites across the United States. Strain names begin with the site code referencing their isolation location. Mean annual temperature (MAT) reflects the 30-year average reported by NOAA.

**Table S2.** Descriptive attributes for 24 *Streptomyces* genomes (previously described in (1)). NCBI accession numbers are associated with BioProject PRJNA401484. Strain names reflect sample sites (see Table S1). Clade membership includes the northern-derived (NDR) and southern-derive (SDR) clades, and the remaining four strains belong to independent (IND) lineages. Genome size in Mb. Genome-wide GC content (%). Number of open reading frames (ORFs). Protein-coding orthologous gene clusters (Genes).

| Strain | NCBI Accession | Clade | Size (Mb) | GC (%) | ORFs | Genes |
| --- | --- | --- | --- | --- | --- | --- |
| b62 | SAMN07606143 | SDR | 7.82 | 71.63 | 7073 | 6568 |
| b84 | SAMN07606144 | NDR | 8.71 | 71.49 | 7657 | 7092 |
| b94 | SAMN07606145 | IND | 8.03 | 72.56 | 6939 | 6502 |
| f150 | SAMN07606147 | SDR | 8.06 | 71.60 | 7320 | 6794 |
| f51 | SAMN07606146 | SDR | 7.72 | 71.77 | 6851 | 6405 |
| gb14 | SAMN07606148 | SDR | 7.88 | 71.67 | 6998 | 6489 |
| man185 | SAMN07606149 | IND | 8.07 | 71.60 | 7244 | 6701 |
| ms115 | SAMN07606150 | SDR | 7.50 | 71.74 | 6776 | 6322 |
| ms184 | SAMN07606151 | SDR | 7.86 | 71.68 | 6958 | 6491 |
| or20 | SAMN07606153 | NDR | 9.08 | 71.54 | 8087 | 7474 |
| or3 | SAMN07606152 | NDR | 8.93 | 71.70 | 7956 | 7329 |
| rh195 | SAMN07606155 | NDR | 8.63 | 71.49 | 7804 | 7245 |
| rh206 | SAMN07606156 | NDR | 8.62 | 71.48 | 7817 | 7262 |
| rh207 | SAMN07606175 | NDR | 8.62 | 71.48 | 7825 | 7268 |
| rh34 | SAMN07606154 | NDR | 8.22 | 71.54 | 7392 | 6897 |
| sk226 | SAMN07606158 | IND | 7.96 | 72.51 | 6930 | 6505 |
| st115 | SAMN07606160 | IND | 8.35 | 71.75 | 7498 | 6966 |
| st140 | SAMN07606161 | SDR | 7.96 | 71.50 | 7184 | 6635 |
| st170 | SAMN07606162 | SDR | 8.08 | 71.42 | 7351 | 6801 |
| st77 | SAMN07606159 | SDR | 8.15 | 71.54 | 7346 | 6784 |
| t99 | SAMN07606163 | SDR | 7.78 | 71.67 | 7076 | 6607 |
| wa1002 | SAMN07606164 | NDR | 8.63 | 71.48 | 7577 | 7038 |
| wa1063 | SAMN07606165 | NDR | 8.84 | 71.39 | 7879 | 7351 |
| wa1071 | SAMN07606166 | NDR | 8.74 | 71.39 | 7755 | 7208 |

**Table S3.** Linear model summary. Table reports coefficient, standard error, t statistic, and *P*-value for explanatory variables. See Figure 2.

|  | **B** | **S.E.** | ***t*** | ***P*-value** |
| --- | --- | --- | --- | --- |
| (Constant) | -30696 | 2022 | -15.18 | < 0.001 |
| ANI | 37173 | 2066 | 17.99 | < 0.001 |
| Clade | 21220 | 3370 | 6.30 | < 0.001 |
| ANI x Clade | -21906 | 3448 | -6.35 | < 0.001 |

**$R^2 = 0.82$ , Adjusted $R^2 = 0.81$**

**Table S4.** Pangenome frequency distributions for NDR and SDR clades. Table reports the number of genes (n) and the proportion (prop) of the total pangenome across frequencies for NDR and SDR. Frequency refers to the number of genomes a gene is present in, ranging from 1–10. Gene pools are categorized by gene frequencies. For example, intermediate-low genes are present in 3– 4 strains, and intermediate-high genes are present in 6–8 strains. *P*-values are from a two proportion z-test, with Bonferrori adjustment for multiple comparisons, evaluating the null hypothesis that the proportion of genes at each frequency is the same for NDR and SDR. Significant *P*-values (< 0.0001) are in bold italics.

| Gene Pool | Frequency | NDR |  | SDR |  | *P*-value |
| --- | --- | --- | --- | --- | --- | --- |
|  |  | n | prop | n | prop |  |
| Unique | 1 | 4132 | 0.302 | 4188 | 0.342 | ***9.8E-11*** |
| Rare | 2 | 926 | 0.068 | 1097 | 0.089 | ***7.3E-10*** |
| Intermediate -<br>*Low* | 3 | 1182 | 0.086 | 494 | 0.040 | ***3.3E-50*** |
|  | 4 | 881 | 0.064 | 377 | 0.031 | ***3.3E-35*** |
|  | 5 | 533 | 0.039 | 260 | 0.021 | ***1.5E-15*** |
| Intermediate -<br>*High* | 6 | 430 | 0.031 | 294 | 0.024 | 0.0032 |
|  | 7 | 230 | 0.017 | 289 | 0.024 | 0.0012 |
|  | 8 | 279 | 0.020 | 303 | 0.025 | 0.21 |
| Near Core | 9 | 854 | 0.062 | 557 | 0.045 | ***2.0E-08*** |
| Core | 10 | 4234 | 0.309 | 4400 | 0.359 | ***3.7E-16*** |

**Figure S1.**
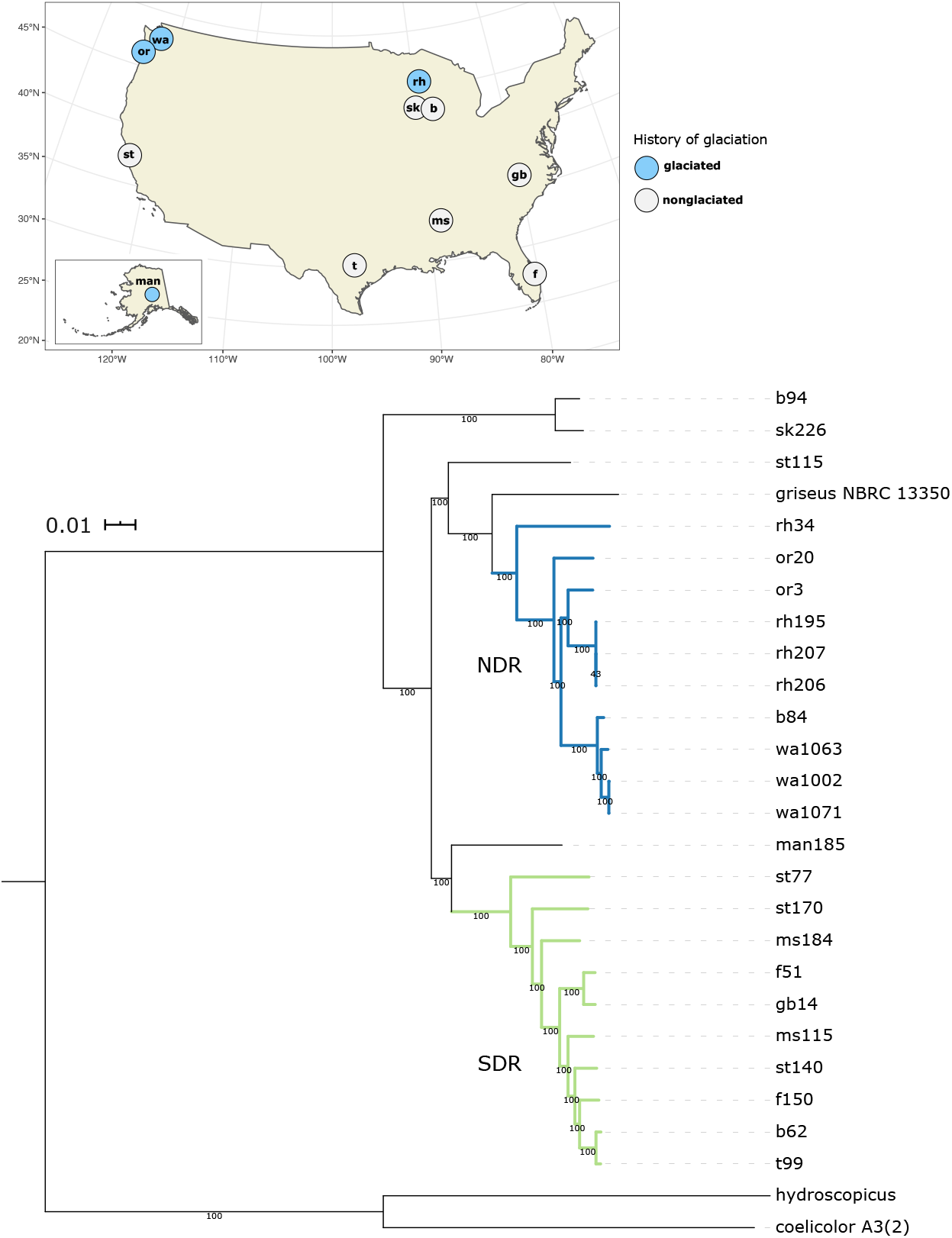
Whole genome phylogeny. *Streptomyces* northern-derived (NDR) and southern-derived (SDR) clades are recently diverged and originate from discrete latitudinal geographic ranges. Phylogenetic relationships were reconstructed from whole genome alignments using maximum likelihood and a GTRGAMMA model of evolution. The scale bar indicates nucleotide substitution per site, and nodes are labeled with bootstrap values. Strains are named according to their sample location (see Table S1). NDR clade branches are blue, and SDR clade branches are green. *Streptomyces griseus* NBRC 13350 is included as the closest taxonomic neighbor. The tree was rooted with *Streptomyces hydroscopicus* and *Streptomyces coelicolor* A3(2). Map shows the sample locations which are labeled with site codes (see Table S1) and colored according to the extent of historical glaciation (see (2)).

**Figure S2.**
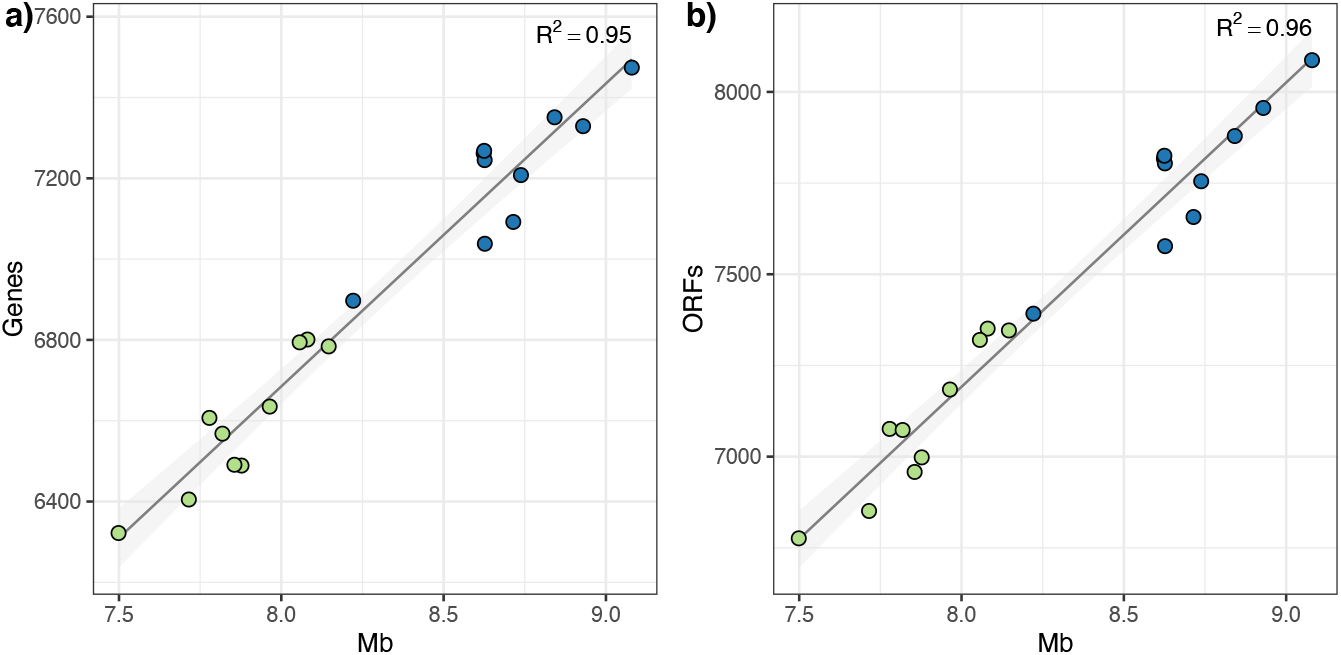
Genome size correlates positively with gene content. Plots show the relationship between genome size in Mb and number of genes (i.e., protein-coding orthologous gene clusters) (a) and open reading frames (ORFs) (b). Circles show values for individual *Streptomyces* genomes (NDR in blue and SDR in green). Lines show the linear regression, and the shaded area is the 95% confidence interval.

**Figure S3.**
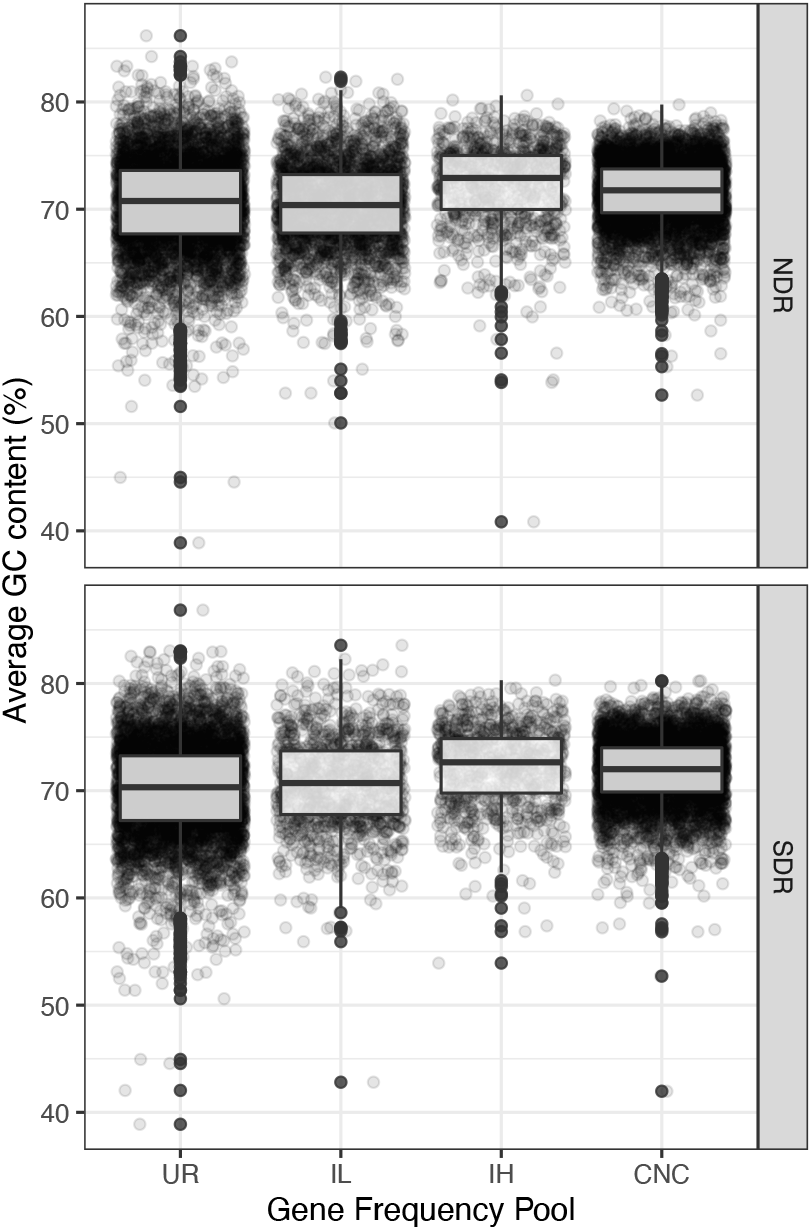
Per-gene GC content. For each gene frequency pool, plots illustrate the distributions of GC skew for NDR (top) and SDR (bottom). Circles show the average GC content (%) for each gene, and boxplots show the medians, interquartile ranges, and 1.5 times interquartile ranges. Gene pools are defined by frequency and include unique-rare (UR) (present in 1–2 strains), intermediate-low (IL) (present in 3–5 strains), intermediate-high (IH) (present in 6–8), and core-near-core (CNC) (present in 9–10 strains) (Table S4).

**Figure S4.**
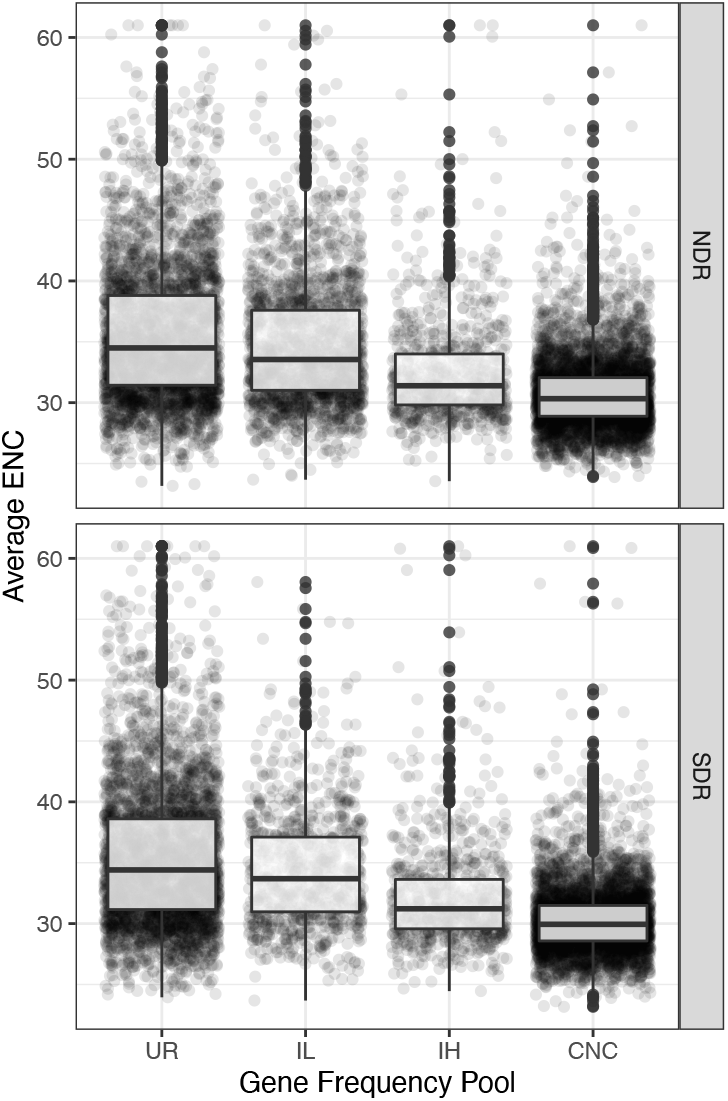
Per-gene codon usage bias. For each gene frequency pool, plots illustrate the distributions of codon usage bias as measured by the effective number of codons (ENC) (3) for NDR (top) and SDR (bottom). Circles show the average ENC for each gene, and boxplots show the medians, interquartile ranges, and 1.5 times interquartile ranges. ENC values range from 61 (indicating no codon usage bias, or all synonymous codons are used in equal frequency), to 2 (indicating extreme codon bias, or extreme favoring of certain codons). Gene pools are defined by frequency and include unique-rare (UR) (present in 1–2 strains), intermediate-low (IL) (present in 3–5 strains), intermediate-high (IH) (present in 6–8), and core-near-core (CNC) (present in 9–10 strains) (Table S4).

**Figure S5.**
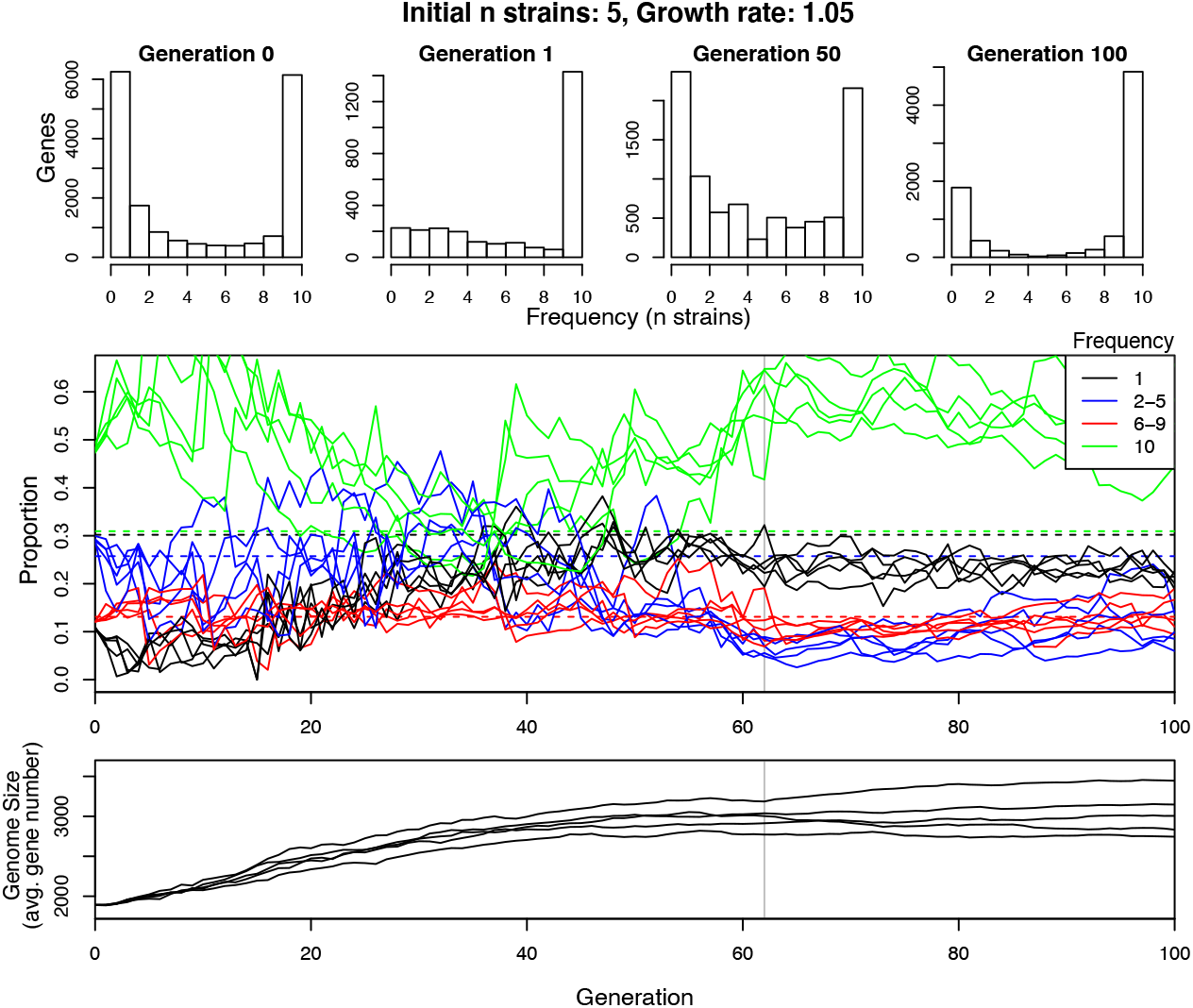
Demographic simulation. We simulated a population range expansion and modeled pangenome dynamics and genome size for 100 generations. *Top:* Pangenome gene frequency distributions from a single randomly selected simulation. Generation 0 shows the frequency distribution of the initial population before the bottleneck. Generation 1 shows the frequency distribution right after the bottleneck. Generations 50 and 100 show the frequency distributions at the subsequent generations. *Middle:* Trajectories of gene frequencies across 100 generations for 5 independent simulations. Colored lines show the proportion of genes within the pangenome present at different frequencies as according to legend. For example, blue lines are the proportions of total genes present in 2–5 strains. Dashed horizontal lines are the observed proportions of the SDR clade (see Table S3). The vertical gray line indicates the generation when the population reached the maximum size. *Bottom:* Genome size. Lines show the average genome size for 5 independent simulations.

## References

1. Vernikos G, Medini D, Riley DR, Tettelin H. Ten years of pan-genome analyses. Curr Opin Microbiol 2015; 23: 148–154.

2. Rasko DA, Rosovitz MJ, Myers GSA, Mongodin EF, Fricke WF, Gajer P, et al. The pangenome structure of Escherichia coli: comparative genomic analysis of E. coli commensal and pathogenic isolates. J Bacteriol 2008; 190: 6881–6893.

3. Reno ML, Held NL, Fields CJ, Burke PV, Whitaker RJ. Biogeography of the Sulfolobus islandicus pan-genome. Proc Natl Acad Sci U S A 2009; 106: 8605–8610.

4. Lefébure T, Bitar PDP, Suzuki H, Stanhope MJ. Evolutionary dynamics of complete Campylobacter pan-genomes and the bacterial species concept. Genome Biol Evol 2010; 2: 646–655.

5. Medini D, Donati C, Tettelin H, Masignani V, Rappuoli R. The microbial pan-genome. Curr Opin Genet Dev 2005; 15: 589–594.

6. Tettelin H, Masignani V, Cieslewicz MJ, Donati C, Medini D, Ward NL, et al. Genome analysis of multiple pathogenic isolates of Streptococcus agalactiae: Implications for the microbial ‘pan-genome’. Proc Natl Acad Sci U S A 2005; 102: 13950–13955.

7. McInerney JO, McNally A, O’Connell MJ. Why prokaryotes have pangenomes. Nature Microbiology 2017; 2: 17040.

8. Brockhurst MA, Harrison E, Hall JPJ, Richards T, McNally A, MacLean C. The ecology and evolution of pangenomes. Curr Biol 2019; 29: R1094–R1103.

9. Azarian T, Huang I-T, Hanage WP. Structure and Dynamics of Bacterial Populations: Pangenome Ecology. In: Tettelin H, Medini D (eds). The Pangenome: Diversity, Dynamics and Evolution of Genomes. 2020. Springer, Cham (CH).

10. Lynch M. Streamlining and simplification of microbial genome architecture. Annu Rev Microbiol 2006; 60: 327–349.

11. Fraser CM, Eisen J, Fleischmann RD, Ketchum KA, Peterson S. Comparative genomics and understanding of microbial biology. Emerg Infect Dis 2000; 6: 505–512.

12. Kuo C-H, Moran NA, Ochman H. The consequences of genetic drift for bacterial genome complexity. Genome Res 2009; 19: 1450–1454.

13. Puigbò P, Lobkovsky AE, Kristensen DM, Wolf YI, Koonin EV. Genomes in turmoil: quantification of genome dynamics in prokaryote supergenomes. BMC Biol 2014; 12: 66.

14. Viklund J, Ettema TJG, Andersson SGE. Independent genome reduction and phylogenetic reclassification of the oceanic SAR11 clade. Mol Biol Evol 2012; 29: 599–615.

15. Giovannoni SJ, Cameron Thrash J, Temperton B. Implications of streamlining theory for microbial ecology. ISME J 2014; 8: 1553–1565.

16. Mira A, Ochman H, Moran NA. Deletional bias and the evolution of bacterial genomes. Trends Genet 2001; 17: 589–596.

17. Moran NA. Microbial minimalism: Genome reduction in bacterial pathogens. Cell 2002; 108: 583–586.

18. McCutcheon JP, Moran NA. Extreme genome reduction in symbiotic bacteria. Nat Rev Microbiol 2011; 10: 13–26.

19. Giovannoni SJ, Tripp HJ, Givan S, Podar M, Vergin KL, Baptista D, et al. Genome streamlining in a cosmopolitan oceanic bacterium. Science 2005; 309: 1242–1245.

20. Brewer TE, Handley KM, Carini P, Gilbert JA, Fierer N. Genome reduction in an abundant and ubiquitous soil bacterium ‘Candidatus Udaeobacter copiosus’. Nat Microbiol 2016; 2: 16198.

21. Konstantinidis KT, Tiedje JM. Trends between gene content and genome size in prokaryotic species with larger genomes. Proc Natl Acad Sci U S A 2004; 101: 3160–3165.

22. Dini-Andreote F, Andreote FD, Araújo WL, Trevors JT, van Elsas JD. Bacterial genomes: habitat specificity and uncharted organisms. Microb Ecol 2012; 64: 1–7.

23. Han K, Li Z-F, Peng R, Zhu L-P, Zhou T, Wang L-G, et al. Extraordinary expansion of a Sorangium cellulosum genome from an alkaline milieu. Sci Rep 2013; 3: 2101.

24. Cordero OX, Hogeweg P. The impact of long-distance horizontal gene transfer on prokaryotic genome size. Proc Natl Acad Sci U S A 2009; 106: 21748–21753.

25. Lerat E, Daubin V, Ochman H, Moran NA. Evolutionary origins of genomic repertoires in bacteria. PLoS Biol 2005; 3: e130.

26. Treangen TJ, Rocha EPC. Horizontal transfer, not duplication, drives the expansion of protein families in prokaryotes. PLoS Genet 2011; 7: e1001284.

27. Bohlin J, Brynildsrud OB, Sekse C, Snipen L. An evolutionary analysis of genome expansion and pathogenicity in Escherichia coli. BMC Genomics 2014; 15: 882.

28. Tsai Y-M, Chang A, Kuo C-H. Horizontal gene acquisitions contributed to genome expansion in insect-Symbiotic Spiroplasma clarkii. Genome Biol Evol 2018; 10: 1526–1532.

29. Wright S. Evolution in Mendelian populations. Genetics 1931; 16: 97–159.

30. Kimura M. Evolutionary rate at the molecular level. Nature 1968; 217: 624–626.

31. Bobay L-M, Ochman H. The Evolution of Bacterial Genome Architecture. Front Genet 2017; 8: 72.

32. Rocha EPC. Neutral theory, microbial practice: Challenges in bacterial population genetics. Mol Biol Evol 2018; 35: 1338–1347.

33. Roselló-Mora R, Amann R. The species concept for prokaryotes. FEMS Microbiol Rev 2001; 25: 39–67.

34. Achtman M, Wagner M. Microbial diversity and the genetic nature of microbial species. Nat Rev Microbiol 2008; 6: 431–440.

35. Shapiro BJ. What microbial population genomics has taught us about speciation. In: Polz MF, Rajora OP (eds). Population Genomics: Microorganisms. 2019. Springer International Publishing, Cham, pp 31–47.

36. Smith NH, Dale J, Inwald J, Palmer S, Gordon SV, Hewinson RG, et al. The population structure of Mycobacterium bovis in Great Britain: clonal expansion. Proc Natl Acad Sci U S A 2003; 100: 15271–15275.

37. Achtman M. Population structure of pathogenic bacteria revisited. Int J Med Microbiol 2004; 294: 67–73.

38. Nübel U, Dordel J, Kurt K, Strommenger B, Westh H, Shukla SK, et al. A timescale for evolution, population expansion, and spatial spread of an emerging clone of methicillin-resistant Staphylococcus aureus. PLoS Pathog 2010; 6: e1000855.

39. Wirth T, Hildebrand F, Allix-Béguec C, Wölbeling F, Kubica T, Kremer K, et al. Origin, spread and demography of the Mycobacterium tuberculosis complex. PLoS Pathog 2008; 4: e1000160.

40. Takuno S, Kado T, Sugino RP, Nakhleh L, Innan H. Population genomics in bacteria: a case study of Staphylococcus aureus. Mol Biol Evol 2012; 29: 797–809.

41. Cornejo OE, Lefebure T, Pavinski 2. Paulina, Lang P, Richards 2. Vincent P., Eilertson K, et al. Evolutionary and population genomics of the cavity causing bacteria Streptococcus mutans. Evolution ; 30: 881–893.

42. Montano V, Didelot X, Foll M, Linz B, Reinhardt R, Suerbaum S, et al. Worldwide population structure, long-term demography, and local adaptation of Helicobacter pylori. Genetics 2015; 200: 947–963.

43. Hewitt G. Some genetic consequences of ice ages, and their role in divergence and speciation. Biological Journal of the Linnean Society 1996; 58: 247–276.

44. Hewitt GM. Genetic consequences of climatic oscillations in the Quaternary. Philos Trans R Soc Lond B Biol Sci 2004; 359: 183–195.

45. Andam CP, Doroghazi JR, Campbell AN, Kelly PJ, Choudoir MJ, Buckley DH. A latitudinal diversity gradient in terrestrial bacteria of the genus Streptomyces. MBio 2016; 7: e02200–15.

46. Choudoir MJ, Doroghazi JR, Buckley DH. Latitude delineates patterns of biogeography in terrestrial Streptomyces. Environ Microbiol 2016; 18: 4931–4945.

47. Choudoir MJ, Buckley DH. Phylogenetic conservatism of thermal traits explains dispersal limitation and genomic differentiation of Streptomyces sister-taxa. ISME J 2018; 12: 2176– 2186.

48. Choudoir MJ, Panke-Buisse K, Andam CP, Buckley DH. Genome surfing as driver of microbial genomic diversity. Trends Microbiol 2017; 25: 624–636.

49. El-Nakeeb MA, Lechevalier HA. Selective isolation of aerobic Actinomycetes. Appl Microbiol 1963; 11: 75–77.

50. Ottow JCG. Rose Bengal as a selective aid in the isolation of fungi and actinomycetes from natural sources. Mycologia 1972; 64: 304.

51. Doroghazi JR, Buckley DH. Widespread homologous recombination within and between Streptomyces species. ISME J 2010; 4: 1136.

52. Kieser, T, Bibb, MJ, Buttner, MJ, Charter, KF, Hopwood, DA. Practical Streptomyces Genetics. 2000. John Innes Foundation, Norwich, UK.

53. Tritt A, Eisen JA, Facciotti MT, Darling AE. An integrated pipeline for de novo assembly of microbial genomes. PLoS One 2012; 7: e42304.

54. Aziz RK, Bartels D, Best AA, DeJongh M, Disz T, Edwards RA, et al. The RAST server: Rapid annotations using subsystems technology. BMC Genomics 2008; 9: 75.

55. Parks DH, Imelfort M, Skennerton CT, Hugenholtz P, Tyson GW. CheckM: assessing the quality of microbial genomes recovered from isolates, single cells, and metagenomes. Genome Res 2015; 25: 1043–1055.

56. Benedict MN, Henriksen JR, Metcalf WW, Whitaker RJ, Price ND. ITEP: an integrated toolkit for exploration of microbial pan-genomes. BMC Genomics 2014; 15: 8.

57. Angiuoli SV, Salzberg SL. Mugsy: fast multiple alignment of closely related whole genomes. Bioinformatics 2011; 27: 334–342.

58. Capella-Gutierrez S, Silla-Martinez JM, Gabaldon T. trimAl: a tool for automated alignment trimming in large-scale phylogenetic analyses. Bioinformatics 2009; 25: 1972– 1973.

59. Tavare S. Some probabilistic and statistical problems in the analysis of DNA sequences. Lectures Math Life Sci 17: 57–86 Warscheid T, Braams J (2000) Biodeterioration of stone: a review. Int Biodeterior Biodegradation 1986; 46: 343–368.

60. Stamatakis A. RAxML-VI-HPC: maximum likelihood-based phylogenetic analyses with thousands of taxa and mixed models. Bioinformatics 2006; 22: 2688–2690.

61. Stamatakis A, Hoover P, Rougemont J, Renner S. A rapid bootstrap algorithm for the RAxML web servers. Syst Biol 2008; 57: 758–771.

62. Schloss PD, Westcott SL, Ryabin T, Hall JR, Hartmann M, Hollister EB, et al. Introducing mothur: Open-source, platform-independent, community-supported software for describing and comparing microbial communities. Appl Environ Microbiol 2009; 75: 7537–7541.

63. Pagès HA, Gentleman P, DebRoy R. Biostrings: Efficient manipulation of biological strings. R package version 2.59. 2020.

64. Elek A, Kuzman M, Vlahovicek K. coRdon: codon usage analysis and prediction of gene expressivity. R package version 1.8.0. 2019.

65. Katoh K, Standley DM. MAFFT Multiple sequence alignment software version 7: Improvements in performance and usability. Mol Biol Evol 2013; 30: 772–780.

66. Talavera, Gerard, Castresana, Jose. Improvement of phylogenies after removing divergent and ambiguously aligned blocks from protein sequence alignments. Syst Biol 2007; 56: 564–577.

67. Suyama M, Torrents D, Bork P. PAL2NAL: robust conversion of protein sequence alignments into the corresponding codon alignments. Nucleic Acids Res 2006; 34: W609– 12.

68. Korber BT. HIV Signature and Sequence Variation Analysis. In: Allen G. Rodrigo and Gerald H. Learn (ed). Computational Analysis of HIV Molecular Sequences. 2000. Dordrecht, Netherlands: Kluwer Academic Publishers, p Chapter 4, pages 55–72.

69. Marttinen P, Croucher NJ, Gutmann MU, Corander J, Hanage WP. Recombination produces coherent bacterial species clusters in both core and accessory genomes. Microbial Genomics 2015; 1.

70. Kim M, Oh H-S, Park S-C, Chun J. Towards a taxonomic coherence between average nucleotide identity and 16S rRNA gene sequence similarity for species demarcation of prokaryotes. Int J Syst Evol Microbiol 2014; 64: 346–351.

71. Ciufo S, Kannan S, Sharma S, Badretdin A, Clark K, Turner S, et al. Using average nucleotide identity to improve taxonomic assignments in prokaryotic genomes at the NCBI. Int J Syst Evol Microbiol 2018; 68: 2386–2392.

72. Haegeman B, Weitz JS. A neutral theory of genome evolution and the frequency distribution of genes. BMC Genomics 2012; 13: 1.

73. Wright F. The ‘effective number of codons’ used in a gene. Gene 1990; 87: 23–29.

74. Slatkin M, Excoffier L. Serial founder effects during range expansion: a spatial analog of genetic drift. Genetics 2012; 191: 171–181.

75. Kimura M. The Neutral Theory of Molecular Evolution. 1983. Cambridge University Press.

76. Edmonds CA, Lillie AS, Cavalli-Sforza LL. Mutations arising in the wave front of an expanding population. Proc Natl Acad Sci U S A 2004; 101: 975–979.

77. Travis JMJ, Münkemüller T, Burton OJ, Best A, Dytham C, Johst K. Deleterious mutations can surf to high densities on the wave front of an expanding population. Mol Biol Evol 2007; 24: 2334–2343.

78. Chuang A, Peterson CR. Expanding population edges: theories, traits, and trade7offs. Glob Chang Biol 2016; 2: 494–512.

79. Doroghazi JR, Buckley DH. Intraspecies comparison of Streptomyces pratensis genomes reveals high levels of recombination and gene conservation between strains of disparate geographic origin. BMC Genomics 2014; 15: 970.

80. Cheng K, Rong X, Huang Y. Widespread interspecies homologous recombination reveals reticulate evolution within the genus Streptomyces. Mol Phylogenet Evol 2016; 102: 246– 254.

81. Andam CP, Choudoir MJ, Vinh Nguyen A, Sol Park H, Buckley DH. Contributions of ancestral inter-species recombination to the genetic diversity of extant Streptomyces lineages. ISME J 2016; 10: 1731–1741.

82. Currat M, Ruedi M, Petit RJ, Excoffier L. The hidden side of invasions: massive introgression by local genes. Evolution 2008; 62: 1908–1920.

83. Gogarten JP, Townsend JP. Horizontal gene transfer, genome innovation and evolution. Nat Rev Microbiol 2005; 3: 679–687.

84. Excoffier L, Foll M, Petit RJ. Genetic consequences of range expansions. Annu Rev Ecol Evol Syst 2009; 40: 481–501.

85. Hildebrand F, Meyer A, Eyre-Walker A. Evidence of selection upon genomic GC-content in bacteria. PLoS Genet 2010; 6: e1001107.

86. Raghavan R, Kelkar YD, Ochman H. A selective force favoring increased G+C content in bacterial genes. Proc Natl Acad Sci U S A 2012; 109: 14504–14507.

87. Bohlin J. Genome expansion in bacteria: the curios case of Chlamydia trachomatis. BMC Res Notes 2015; 8: 512.

88. Hallatschek O, Hersen P, Ramanathan S, Nelson DR. Genetic drift at expanding frontiers promotes gene segregation. Proc Natl Acad Sci U S A 2007; 104: 19926–19930.

89. Moran NA. Accelerated evolution and Muller’s rachet in endosymbiotic bacteria. Proc Natl Acad Sci U S A 1996; 93: 2873–2878.

90. Rispe C, Moran NA. Accumulation of deleterious mutations in endosymbionts: Muller’s ratchet with two levels of selection. Am Nat 2000; 156: 425–441.

91. Kuo C-H, Ochman H. The fate of new bacterial genes. FEMS Microbiol Rev 2009; 33: 38– 43.

92. Lobkovsky AE, Wolf YI, Koonin EV. Gene frequency distributions reject a neutral model of genome evolution. Genome Biol Evol 2013; 5: 233–242.

93. Domingo-Sananes MR, McInerney JO. Selection-based model of prokaryote pangenomes. bioRxiv. 2019; doi:10.1101/782573.

94. Cordero OX, Ventouras L-A, DeLong EF, Polz MF. Public good dynamics drive evolution of iron acquisition strategies in natural bacterioplankton populations. Proc Natl Acad Sci U S A 2012; 109: 20059–20064.

95. McNally A, Kallonen T, Connor C, Abudahab K, Aanensen DM, Horner C, et al. Diversification of colonization factors in a multidrug-resistant Escherichia coli lineage evolving under negative frequency-dependent selection. MBio 2019; 10: e00644–19.

96. Mira A, Klasson L, Andersson SGE. Microbial genome evolution: sources of variability. Curr Opin Microbiol 2002; 5: 506–512.

97. Mira A, Martín-Cuadrado AB, D’Auria G, Rodríguez-Valera F. The bacterial pan-genome: a new paradigm in microbiology. Int Microbiol 2010; 13: 45–57.

98. Boucher Y, Doolittle WF. The role of lateral gene transfer in the evolution of isoprenoid biosynthesis pathways. Mol Microbiol 2000; 37: 703–716.

99. Pál C, Papp B, Lercher MJ. Adaptive evolution of bacterial metabolic networks by horizontal gene transfer. Nat Genet 2005; 37: 1372–1375.

100. Gould SJ, Lewontin RC. The spandrels of San Marco and the Panglossian paradigm: a critique of the adaptationist programme. Proc R Soc Lond B Biol Sci 1979; 205: 581–598.

## Supplementary References

101. Choudoir MJ, Buckley DH. 2018. Phylogenetic conservatism of thermal traits explains dispersal limitation and genomic differentiation of Streptomyces sister-taxa. ISME J 12:2176–2186.

102. Andam CP, Doroghazi JR, Campbell AN, Kelly PJ, Choudoir MJ, Buckley DH. 2016. A Latitudinal Diversity Gradient in Terrestrial Bacteria of the Genus Streptomyces. MBio 7:e02200–15.

103. Wright F. 1990. The “effective number of codons” used in a gene. Gene 87:23–29.

